# Evolution of a minimal cell

**DOI:** 10.1101/2021.06.30.450565

**Authors:** RZ Moger-Reischer, JI Glass, KS Wise, L Sun, D Bittencourt, M Lynch, JT Lennon

**Affiliations:** Department of Biology, Indiana University, Bloomington, IN 47405, USA; J. Craig Venter Institute, La Jolla, CA 92037, USA; Sorrento Therapeutics, Inc., San Diego, CA 92121 USA; Embrapa Genetic Resources and Biotechnology, Brasília, 70770-917, Brazil; Arizona State University, Tempe, AZ 85287, USA

## Abstract

Possessing only essential genes, a minimal cell can reveal mechanisms and processes that are critical for the persistence and stability of life. Here, we report on how a synthetically constructed minimal cell contends with the forces of evolution compared to a non-minimized cell from which it was derived. Genome streamlining was costly, but 80% of fitness was regained in 2000 generations. Although selection acted upon divergent sets of mutations, the rates of adaptation in the minimal and non-minimal cell were equivalent. The only apparent constraint of minimization involved epistatic interactions that inhibited the evolution of cell size. Together, our findings demonstrate the power of natural selection to rapidly optimize fitness in the simplest autonomous organism, with implications for the evolution of cellular complexity.

## INTRODUCTION

The complexity of an organism is reflected in the number of genes it possesses, a quantity that varies by orders of magnitude across the tree of life. While some obligately endosymbiotic bacteria have fewer than 200 protein-coding genes, many plant and animal genomes contain more than 20,000 genes (Lynch 2007). In principle, the simplest organism would have no functional redundancies and possess only the minimum number of genes essential for life. Any mutation in such an organism could lethally disrupt one or more cellular functions, placing constraints on evolution, as reflected in the fact that essential proteins change more slowly than those encoded by dispensable genes (Graur and Li 2000; Hahn and Kern 2005). Furthermore, organisms with streamlined genomes have fewer targets upon which positive selection can act, thus limiting opportunities for adaptation. Understanding how organisms with simplified genomes overcome these evolutionary challenges has important implications for long-standing problems in biology, including the treatment of clinical pathogens, the persistence of host-associated endosymbionts, the refinement of engineered microorganisms, and the origin of life itself (Leprince et al. 2012; Moran and Bennett 2014; Glass et al. 2017).

The cell is the simplest independent functional unit of life. Yet even unicellular model organisms that are touted for their tractability are complex, possessing thousands of genes and proteins, many of which remain uncharacterized even after decades of in-depth interrogation. The quest for the simplest organism has been aided by advances in synthetic biology, which involves the redesign or novel construction of biological parts and modules (Benner and Sismour 2005; Glass et al. 2017). Promoted as a way to address practical challenges for society (Cameron et al. 2014; DeLisi 2019), synthetic biology also provides a platform for developing powerful simplest-case models through streamlining, whereby nonessential chromosomal sequences are removed from an organism’s genome (Leprince et al. 2012; Hutchison et al. 2016; Glass et al. 2017; Richardson et al. 2017; Lachance et al. 2019). Guided by such strategies, a “minimal cell” was constructed with a genome containing only the smallest set of genes required for autonomous cellular life (Hutchison et al. 2016; Breuer et al. 2019). While these efforts succeeded in experimentally identifying the genetic requirements for basic cellular processes such as metabolism and cell division, it remains unclear how a minimal cell should respond to the forces of evolution, particularly given the limited raw materials upon which natural selection can operate as well as the uncharacterized input of new mutations.

To gain insight into the dynamics and outcomes of evolution in a minimal cell, we conducted experiments with synthetic strains of *Mycoplasma mycoides* (Hutchison et al. 2016; Breuer et al. 2019). The minimal cell, *M. mycoides* JCVI-syn3B, was derived from a non-minimal strain, *M. mycoides* JCVI-syn1.0, by reducing the chromosome from 901 to 493 genes, resulting in the smallest genome of any known free-living organism (Hutchison et al. 2016; Breuer et al. 2019). With these two strains, we first investigated whether genome minimization, which included the removal of two DNA replication genes and eight DNA repair genes, altered the rate and spectrum of new mutations in the minimal cell relative the non-minimal organism. Second, with knowledge of the mutational input, we evaluated whether genome minimization altered the rate and mechanisms of evolution in response to natural selection, as measured using whole-genome sequencing, estimates of population fitness, and phenotypic changes in cell size.

## RESULTS AND DISCUSSION

### Highest recorded mutation rate is robust to genome minimization

Through serial bottlenecking under relaxed selection, we conducted mutation-accumulation experiments with populations of *Mycoplasma mycoides* (see Methods). The number of mutations per nucleotide per generation for the non-minimal cell (3.13 ± 0.12 × 10^-8^, mean ± SEM) was indistinguishable from that of the minimal cell (3.25 ± 0.16 × 10^-8^) (*t*_140_ = 0.43, *P* = 0.667, Fig. 1a). These mutation rates, which are the highest recorded for any cellular organism, are consistent with other reports where organisms with smaller genomes have higher mutation rates (Kuo and Ochman 2009; Sung et al. 2012; Lynch et al. 2016; Long et al. 2018). For example, *Mesoplasma florum* also has an exceptionally high mutation rate (1.16 ± 0.07 × 10^-8^) (Sung et al. 2012), which may reflect the fact that these closely related bacteria both lack mismatch repair, a system for recognizing and fixing errors that are made during DNA replication (Carvalho et al. 2005). Our data are also consistent with predictions from the drift-barrier hypothesis. This theory posits that mutation rates evolve downward until the selective advantage of another incremental decrease in the mutation rate is small enough to be effectively neutral and outweighed by genetic drift (Sung et al. 2012; Lynch et al. 2016). In other words, populations with lower effective population size (*N*_e_) experience stronger drift, and hence evolve higher mutation rates (Lynch et al. 2016). Notably, the natural history (obligate pathogen) and genomic features (small genome size and low GC content) of *M. mycoides* and *Me. florum* suggest these mollicute bacteria most likely have low *N*_e_ (Moran 2003; Kuo and Ochman 2009; Hershberg 2015; Long et al. 2018).

**Fig. 1.**
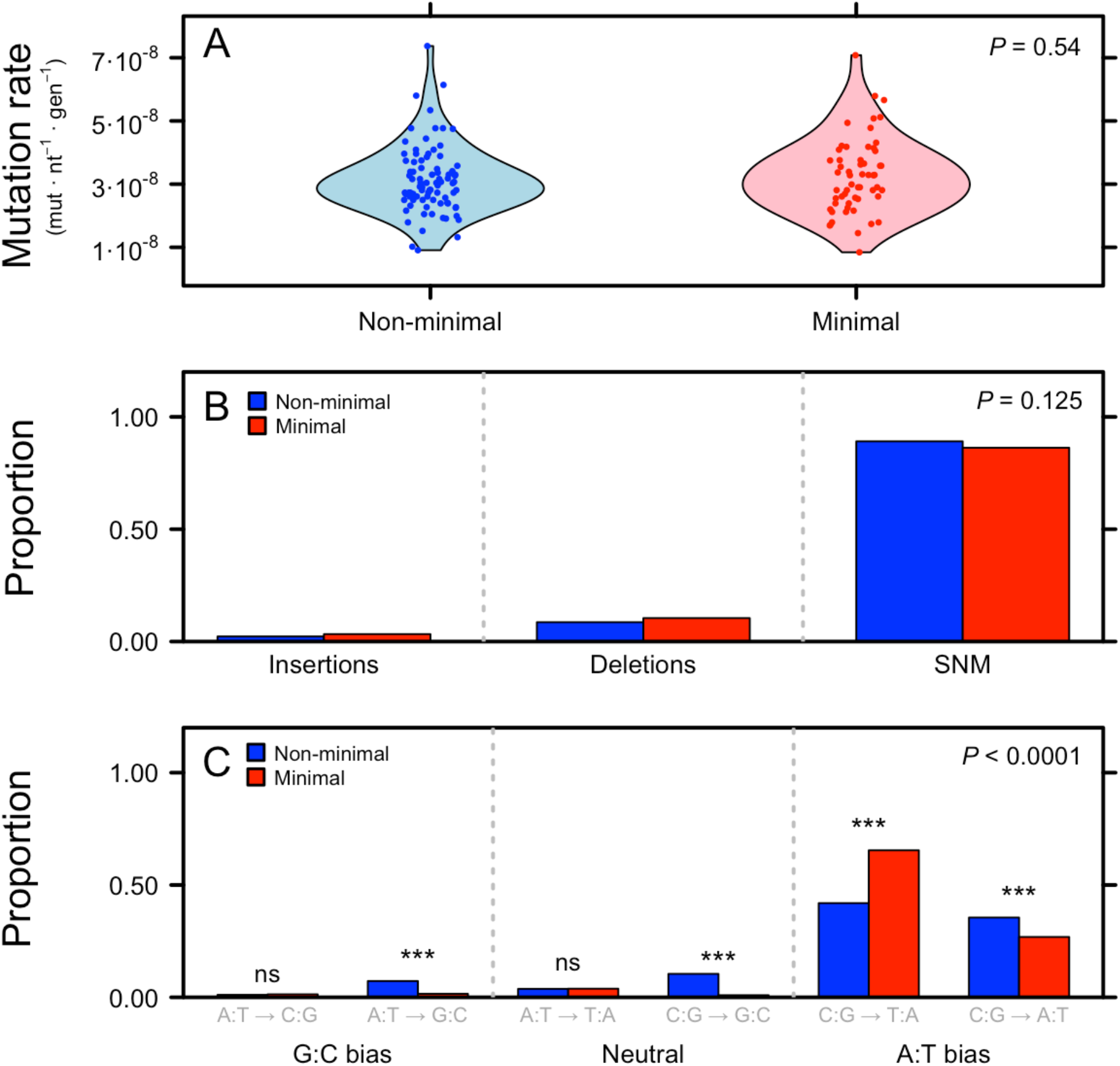
Mutation rate and spectrum of the minimal and non-minimal cell estimated from mutation accumulation experiments. (**A**) Although *M. mycoides* has the highest recorded mutation rate (base substitutions and indels), it was not affected by genome minimization. (**B)** The proportions of insertions, deletions, and single nucleotide mutations (SNMs) were also the same for the minimal and non-minimal cell. (**C**) Among SNMs, which accounted for 88% of all mutations, the minimal cell exhibited a stronger AT bias in its mutation spectrum than the non-minimal cell, particularly in the C:G-to-A:T category. In this panel, ns = not significant and *** = *P* < 0.001.

### Small effect of genome minimization on mutational spectrum

Although the mutation rate was unaffected by minimization, the types of mutations that arise in a population can still influence evolution. Therefore, we compared the spectra of mutations in the minimal and non-minimal *M. mycoides.* Overall, the composition of mutation types (insertions, deletions, and single-nucleotide mutations) was unaffected by genome minimization (*χ*^2^_2_ = 4.16, *P* = 0.125) (Fig. 1b). Both strains exhibited bias toward deletions over insertions (non-minimal: *χ*^2^_1_ = 21.3, *P* = 4.0 × 10^-6^; minimal: *χ*^2^_1_ = 68.4, *P* < 2.2 × 10^-16^), consistent with findings reported for other bacteria (Kuo and Ochman 2009; Hershberg 2015; Sung et al. 2016). However, the composition of single-nucleotide mutations (SNMs), which constituted the largest category of mutations (88%), differed between the minimal and non-minimal cell (Monte Carlo *χ*^2^ = 69.9, *P* = 1.0 × 10^-4^). Mutations from a G or C nucleotide to an A or T nucleotide occurred at a rate ~30-fold higher in the non-minimal cell (*χ*^2^_1_ = 3736, *P* < 2.2 × 10^-16^) and ~100-fold higher in the minimal cell (*χ*^2^_1_ = 1444, *P* < 2.2 × 10^-16^) compared to mutations from A or T to G or C (Fig. 1c). As a consequence, *M. mycoides* exhibits more A:T bias than any known organism (Hershberg 2015; Long et al. 2018).

There are a few explanations for the extreme A:T mutational bias in the minimal cell. First, *M. mycoides* naturally lacks Dut, a dUTPase protein that draws down the intracellular concentration of dUTP, thereby avoiding its misincorporation into DNA (Tye and Lehman 1977). Because adenine would pair with a misincorporated uracil, the lack of Dut should lead to an increased frequency of A:T-biased mutations in the spectrum of cells without Dut (Sedwick et al. 1986). However, since both cells lack Dut, this mechanism is insufficient to explain the discrepancy in A:T bias between the non-minimal and minimal cells, respectively (*χ*^2^_1_ = 21.8, *P* = 3.08 × 10^-6^). Instead, the difference in A:T bias is more likely due to the removal of *ung* during minimization (Hutchison et al. 2016). This nonessential DNA repair gene encodes for the uracil-DNA glycosylase enzyme Ung, which excises ectopic uracils from DNA via hydrolysis. The process results in a free uracil and an abasic site in the DNA strand (Lindahl et al. 1977), which allows the correct nucleotide to be inserted using base complementarity. Without Ung, the uracil would remain in place and be paired with A during DNA replication, even if the original nucleotide was not T. One common cause for the presence of uracil in DNA is the deamination of cytosine (Lindahl 1974). Thus, a cell lacking *ung* would be enriched in C-to-T mutations when C is replaced by U via deamination and U is replaced by T in subsequent DNA replication via base-pairing with A. In the minimal cell, we observed an excess of these C:G-to-T:A mutations relative to the non-minimal bacterium (*χ*^2^_1_ = 89.8, *P* = 1.60 × 10^-20^), consistent with this aspect of the mutation spectrum being related to the minimal cell lacking *ung* (Fig. 1c).

### Rapid recovery of fitness in the minimal cell

We next considered how populations of the minimal and non-minimal cell would respond to natural selection. With measured mutation rates of ~3 × 10^-8^ per nucleotide per generation and population sizes in excess of 10^7^ individuals, we estimated that a new mutation would hit every nucleotide in the genome >250 times during 2000 generations of experimental evolution. Thus, given their equal mutation rates per site, populations of either cell type should not be limited by the availability of genetic variation to fuel adaptation, suggesting that any differences in the ways the two strains adapt would be driven by alterations in genome architecture created by minimization.

To study natural selection, we passaged replicate populations of *M. mycoides* for ~2000 generations (see Methods), a period during which rapid adaptation is often observed (Vasi et al. 1994; Gifford et al. 2011). At the end of the experiment, we measured the fitness (competitive growth rates) of all ancestors and evolved populations (see Methods). For the ancestral strains, we determined that genome minimization led to a 50% reduction in fitness (Fig. 2). Despite this major initial cost, the minimal cell rapidly regained fitness during the course of evolution. Approximately 80% of the fitness lost to minimization was recovered during 300 days of serial passaging (Fig. 2). This suggests that a streamlined *M. mycoides* genome is not inherently crippled and can perform almost as well as the non-minimized cell following readaptation.

**Fig. 2.**
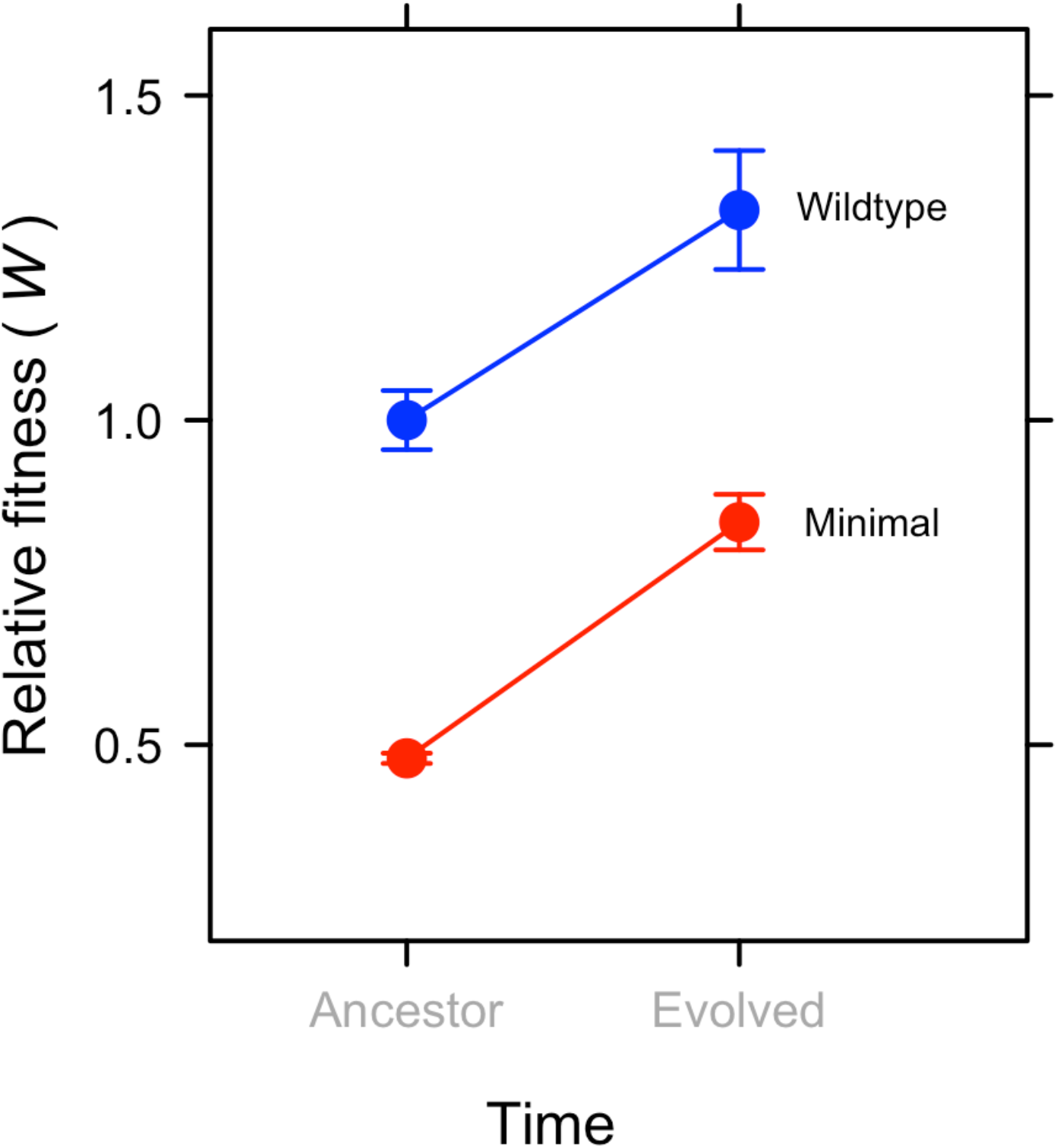
Genome minimization resulted in a 50% reduction in relative fitness, but 80% of this cost was regained over 2000 generations of evolution. Despite removal of nearly half of its genome, the minimal cell adapted at the same rate as the non-minimal cell. Data represent mean ± SEM. Because the experiment was initiated with a single clone, error bars for the ancestral time point were calculated from experimental replicates, while error bars for evolved lines were calculated from replicate populations.

Contrary to our initial predictions, adaptation was not constrained by genome minimization (Fig. 2). The rate of fitness gain was the same for replicate populations of the non-minimal and minimal cells (*t*_6_ = 0.40, *P* = 0.704, Fig. 2). This interpretation was bolstered by results from population-genomic sequencing (see Methods). The relative ratio of nonsynonymous to synonymous fixed SNMs (*d_N_*/*d_S_*) did not differ between the two cell types (*t*_6_ = 1.18, *P* = 0.859, Fig. S1), consistent with the interpretation that the rates of molecular evolution were comparable even though all of the genes in the minimal cell are critical for fitness (Graur and Li 2000; Hahn and Kern 2005).

### Divergent mechanisms of adaptation

Using a combination of statistical simulation and reverse genetics, we identified mutations that likely contributed to the observed patterns of adaptation. First, we analyzed the gene-by-population matrix for non-synonymous mutations that arose in essential genes during the natural selection experiment (see Methods). The two cell types acquired mutations in different sets of essential genes (PERMANOVA, *F*_7_ = 4.12, *P* = 0.029, Fig. 3), suggesting that, despite adapting at similar rates, the populations evolved via divergent routes. To explore this hypothesis, we looked for genes that acquired a higher number of non-synonymous mutations than expected under assumptions of neutrality (see Methods). We identified 16 genes in the non-minimal genome and 14 in the minimal genome that were potential targets of positive selection (Table 1). Second, we used reverse genetics to experimentally verify that one of the common types of mutation observed in replicate populations of both strains was in fact beneficial (Table S1). Using CRISPR editing, we recreated *ftsZ* C-terminal nonsense mutations by inserting an *ftsZ* E315* mutation into the ancestral genomes of the minimized and non-minimized strains (see Methods). Competition assays with the constructs revealed that this putatively adaptive mutation conferred a 25% fitness increase in the non-minimal cell and a 14% fitness increase in the minimal cell (Fig. S2).

**Fig. 3.**
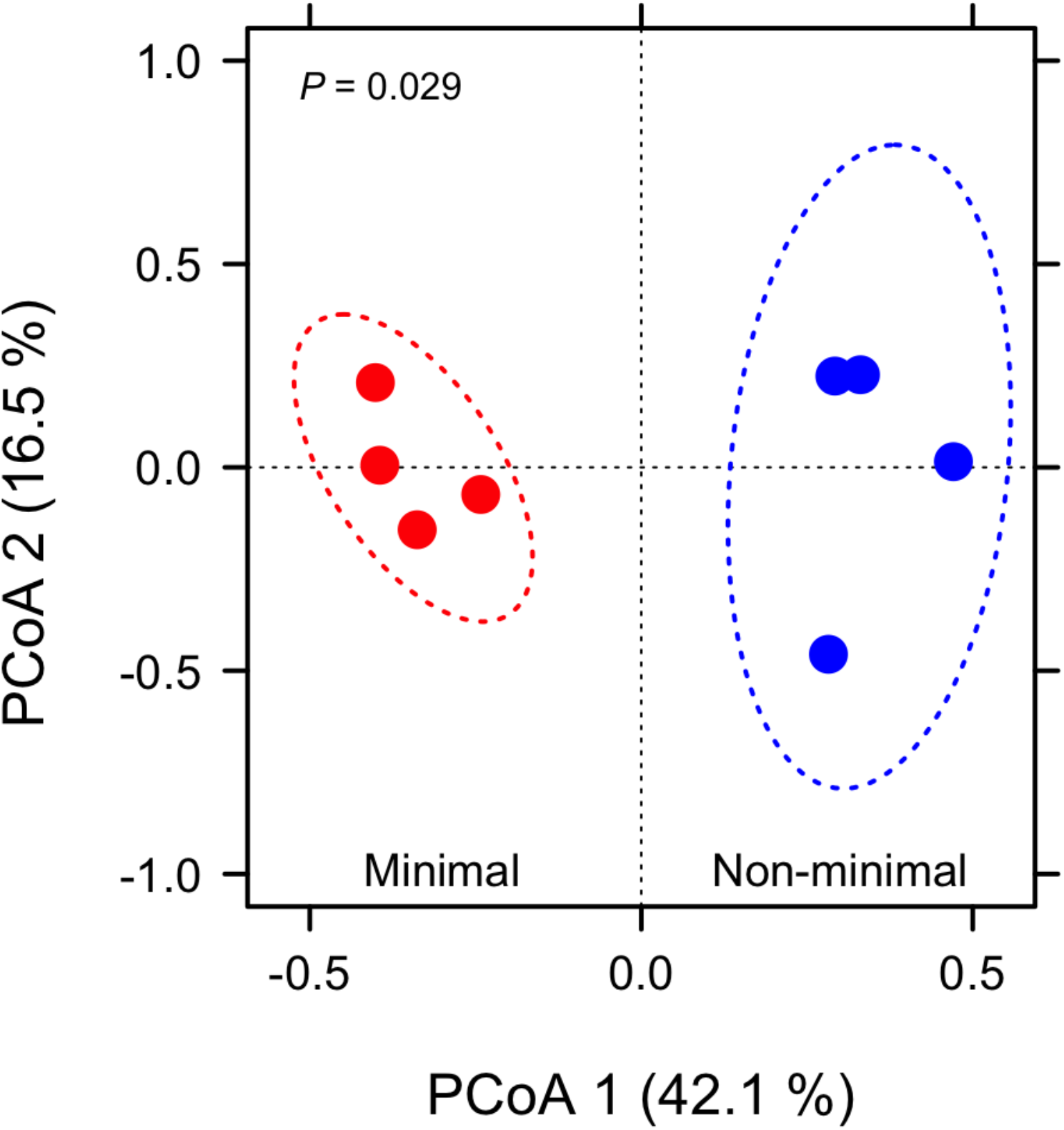
The non-minimal cell and minimal cell acquired adaptive mutations in different sets of shared essential genes (*P* = 0.029) over 2000 generations of evolution (Table 1) as depicted in ordination from a principal coordinates analysis (PCoA) created with a gene-by-population matrix using Bray-Curtis distance metric. Dashed lines represent 95% confidence ellipses around replicate populations (solid symbols).

**Table 1.**
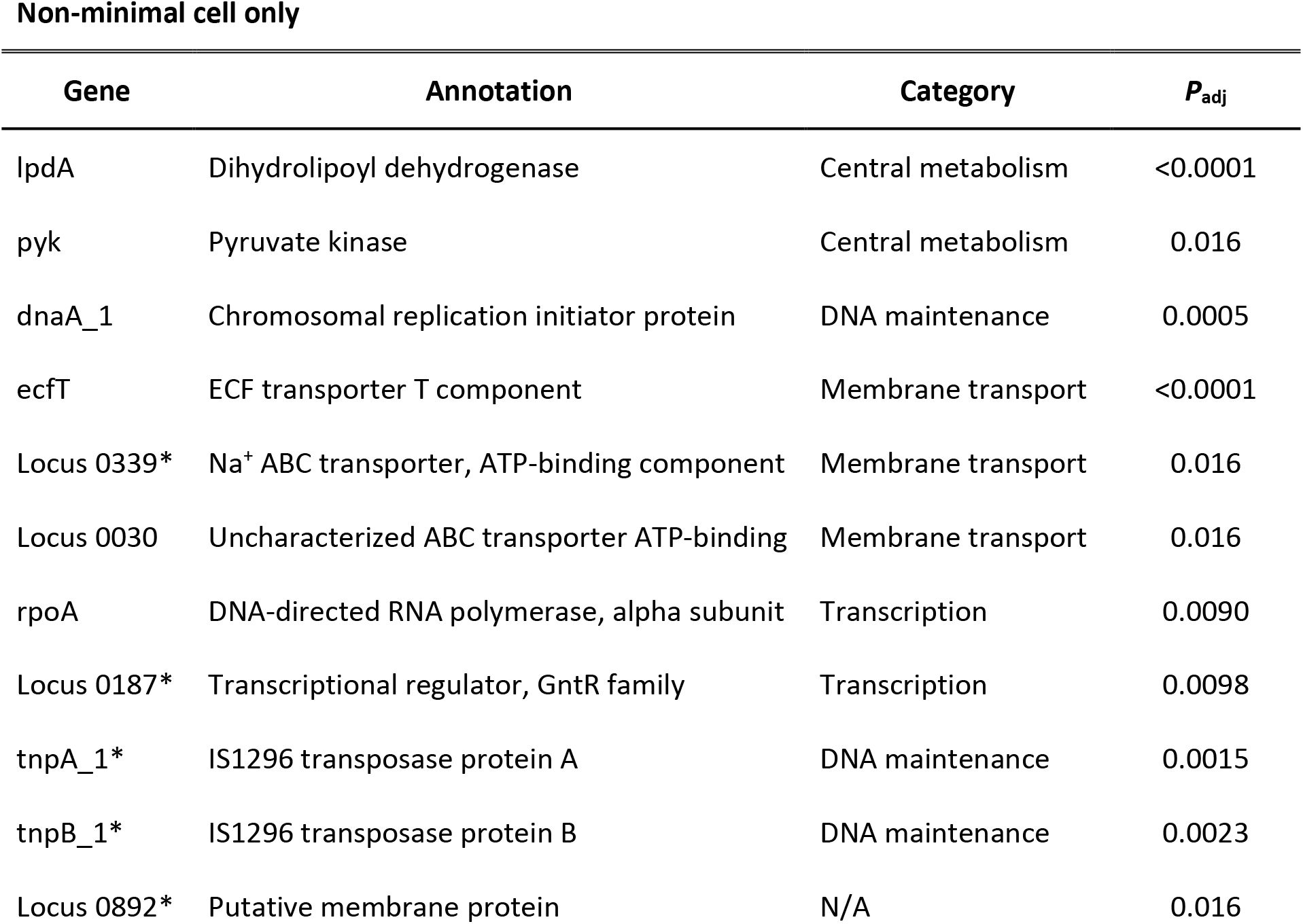
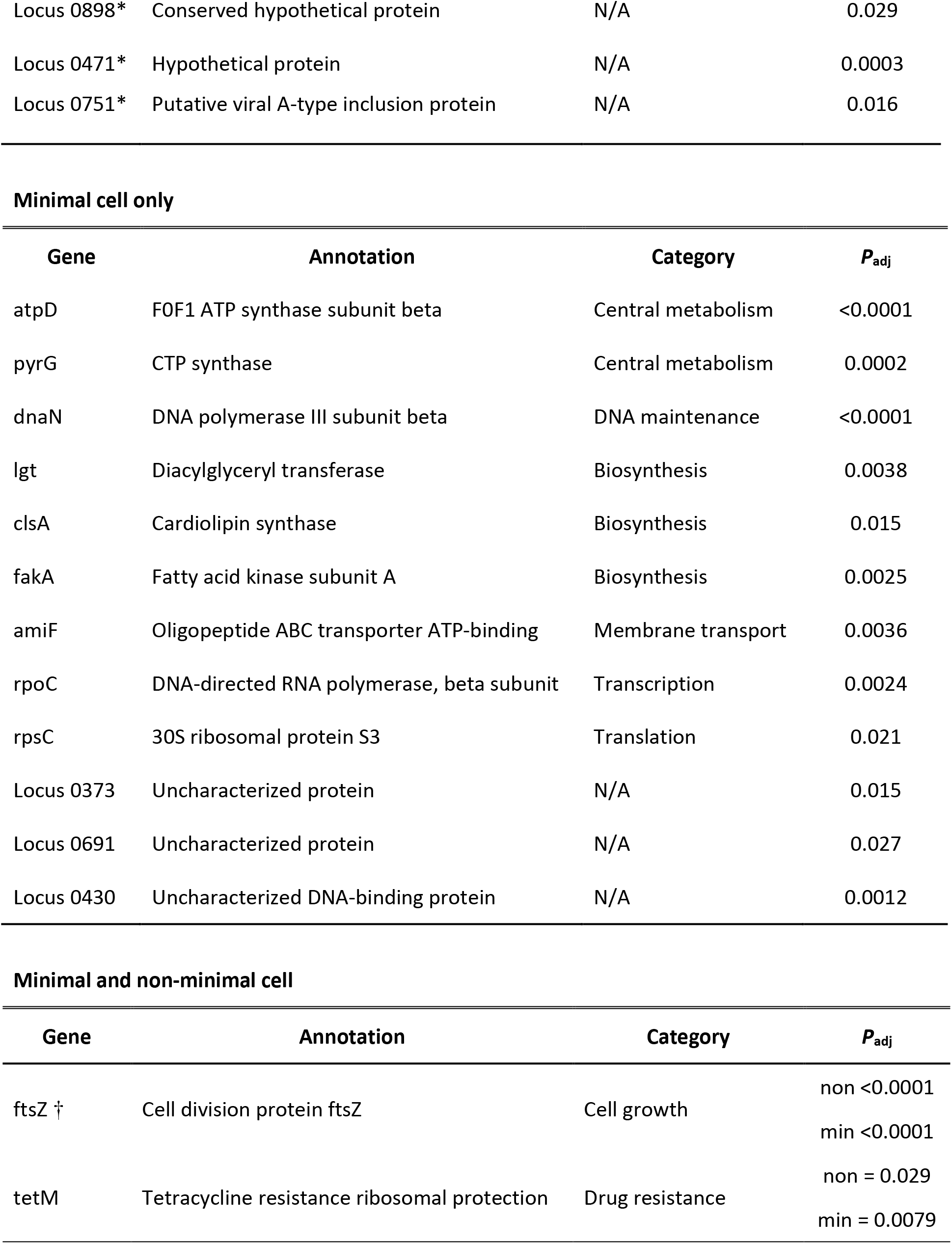
Mutations in genes that were identified as being under positive selection during 2000 generations of experimental evolution. Statistical simulations were performed to find genes acquiring more mutations than expected to occur by chance, indicative of positive selection increasing the allelic proportion of such mutations. *P*_adj_ corresponds to significance following Benjamin-Hochberg correction to account for multiple comparisons. Genes are assigned to categories based on the secondary functional classifications (Breuer et al. 2019). Note that “Central metabolism” corresponds to “Central carbon metabolism” in the original source (Breuer et al. 2019). While most nonessential genes are absent from the minimal cell (*), a few nonessential genes were retained in *M. mycoides* JCVI-syn3B to facilitate cultivation and robust growth (†). Uncharacterized genes did not fall into defined category (N/A). For mutations that were found in both strains, “non” refers to non-minimal and “min” refers to minimal.

Comparative analysis of genes under positive selection provided insight into the functional consequences of adaptation in the minimal cell. We hypothesized that mutations in genes related to membrane transport would be critical for adaptation because the minimal cell relies on the import and export of metabolites and other biomolecules for metabolism (Antczak et al. 2019; Breuer et al. 2019). However, mutations in membrane transport functions were enriched to a similar degree in both cell types over 2000 generations of evolution (Fisher’s Exact Test *P* = 0.934). Instead, we detected a signal of enrichment for mutations in biosynthetic genes for the minimal cell (Fisher’s Exact Test *P* = 0.090), including those involved in lipid metabolism. Specifically, *fakA* and *clsA* (Table 1) are considered essential for synthesizing cardiolipin and other lipids from free fatty acids (Breuer et al. 2019), which are important for the construction of cell membranes and the regulation of cell division. The gene *lgt* is also critical for membrane construction, encoding the protein that transfers diacylglyceryl moieties to anchor surface lipoproteins in the lipid bilayer (Breuer et al. 2019). Thus, metabolic innovations involving lipid synthesis and distribution may be more important for the minimal cell than enhanced acquisition of metabolites that are already present in the growth medium.

### Genome minimization constrains evolution of cell size

The size of single-celled organisms is variable and often linked to fitness in complex ways (Gounand et al. 2016; Mei et al. 2009; Chien et al. 2012). In resource-rich environments, cell size tends to be positively correlated with growth rate, one of the most important components of fitness (Vasi et al. 1994; Mongold and Lenski 1996; Mei et al. 2009; Chien et al. 2012; Hill et al. 2013). For example, in the first 2000 generations of a classic long-term evolution experiment, cell volume and fitness concomitantly increased by 50% and 30%, respectively (Vasi et al. 1994). While an increase in size can accommodate more macromolecules needed for growth and division, it also decreases a cell’s surface-to-volume ratio which increases molecular diffusion times. Given these opposing pressures, we evaluated how cell size changed in response to genome streamlining over the course of evolution. Prior to initiating experiments, we observed that the spherically shaped non-minimal cell was 575 ± 28 nm (mean ± SEM) and the minimal cell was 422 ± 23 nm (see Methods), comparable to measures of cell diameter previously reported (Hutchison et al. 2016; Breuer et al. 2019). Following 2000 generation of evolution, the non-minimal cell exhibited a 73% increase in diameter (993 ± 53 nm, *t*_3_ = 7.84, *P* = 0.004) and a corresponding five-fold increase in volume compared to its ancestor (Fig. 4; Table S2). In contrast, the size of the minimal cell did not change during evolution (*t*_3_ = 0.85, *P* = 0.459).

**Fig. 4.**
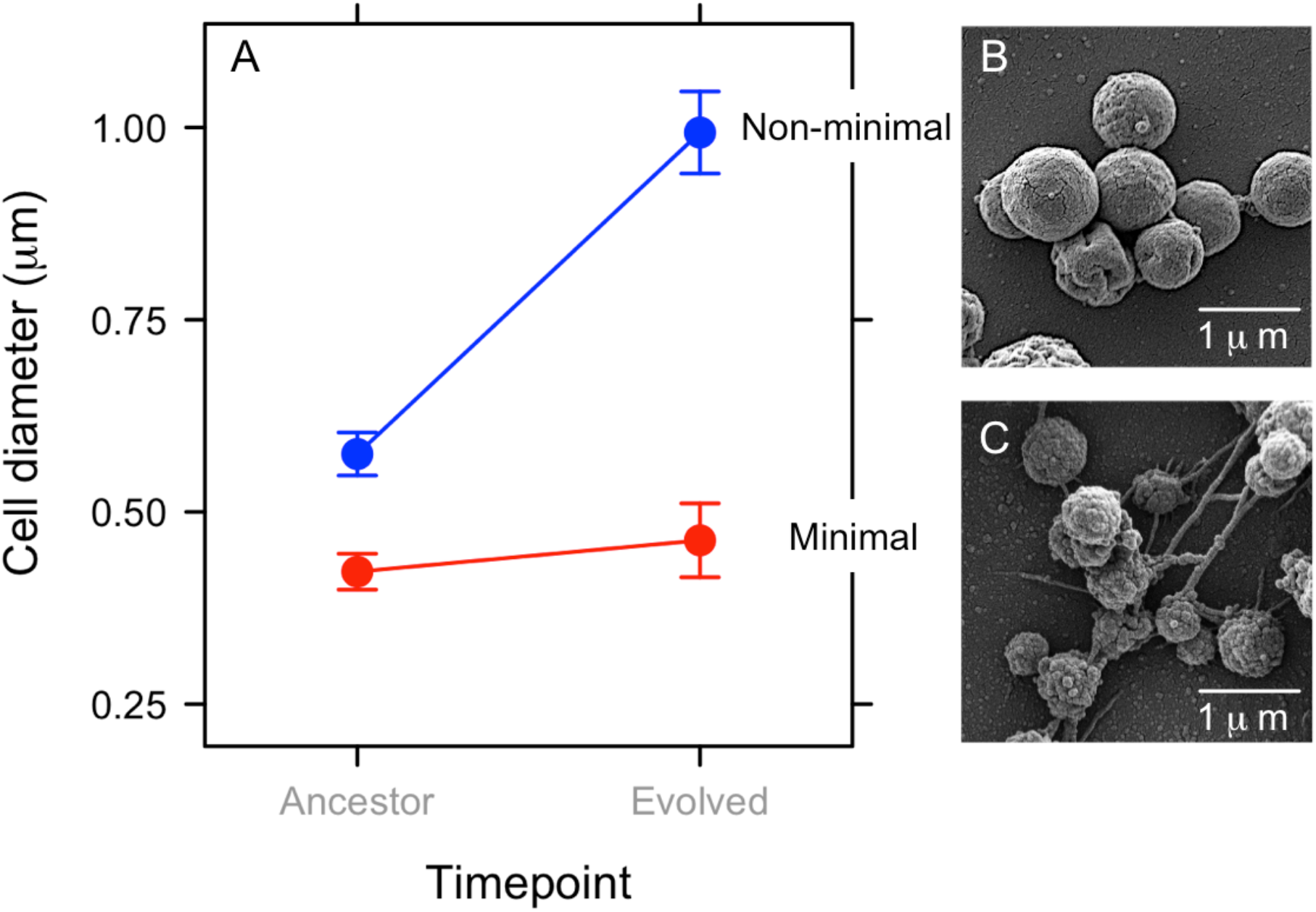
Effect of genome minimization on the evolution of cell size. (**A**) The size of non-minimal cells increased by 73% over 2000 generations of evolution, while the size of minimal cell remained the same. Data represent means ± SEM based on images obtained via phase contrast microscopy. Scanning electron micrographs obtained from evolved replicate populations of the (**B**) non-minimal and (**C**) minimal cell.

While cell size is a complex multigenic trait, previous studies have implicated *ftsZ* with changes in morphology of the minimal cell (Pelletier et al. 2021). It encodes for FtsZ, a protein that localizes to the midcell and determines the site of membrane constriction during cell division. Prevalent among diverse lineages of bacteria and archaea (McQuillen and Xiao 2020; Liao et al. 2021), *ftsZ* is nevertheless nonessential in *M. mycoides*. However, cells lacking *ftsZ* exhibit aberrant cell division and morphology (Hutchison et al. 2016; Breuer et al. 2019; Pelletier et al. 2021). Thus, along with 18 other nonessential genes, *ftsZ* was retained in JCVI-syn3B to aid in culture maintenance and stable growth (Breuer et al. 2019; Pelletier et al. 2021). In our study, *ftsZ* was consistently mutated over 2000 generations of evolution and was identified as a target of positive selection in both the minimal and non-minimal cell (Table 1, Fig. S2). Introduction of this termination codon could eliminate the C-terminal region of FtsZ, known to interact with membrane-associated products that recruit FtsZ (Cohan et al. 2020), The early stop codon could also create a transcriptional polar effect (Graffeuil et al. 2020) that reduces expression of two adjacent down-stream genes within a likely polycistronic operon: MMSYN1_0521, an ortholog of cell division protein *sepF* and MMSYN1_0520, encoding a hydrolase of unknown functions. Irrespective of mechanism, we demonstrated that mutations in *ftsZ* had a non-additive effect that contributed to the evolutionary divergence of cell size. Using scanning electron microscopy, we documented that the *ftsZ* E315* nonsense mutation in the ancestral non-minimal cell led to a 25% increase in cell diameter (*t*_118_ = 5.56, *P* = 8.71 × 10^-8^) and a corresponding two-fold increase in cell volume. In contrast, the same *ftsZ* nonsense mutation in the ancestral minimal cell led to a 19% decrease in the cell diameter (*t*_122_ = 3.29, *P* = 0.001, Fig. S4), which reduced cell volume by half. Thus, the *ftsZ* E315* mutation recapitulated nearly 60% of the evolved divergence in cell size, indicating that FtsZ plays a central role in cell size in *M. mycoides*.

Although changes in *ftsZ* had opposing effects on the size of the minimal and non-minimal cell, mutations in this gene were adaptive for both strains (Fig. S2, Table 1). Smaller cell size may be advantageous for the minimal cell. Owing to its slower growth rate, JCVI-syn3B would have spent more time in an exponential growth phase in our evolution experiment. Some studies have shown that smaller cell size is selected for when bacteria evolve under such conditions (Gounand et al. 2016) even though other studies report a positive association between metabolic rate and body mass across species (DeLong et al. 2010). Meanwhile, the faster-growing non-minimal cell should experience bouts of feast-or-famine over thousands of generations in a serial batch environment. Under such conditions, repeated transitions between exponential and stationary growth phases has been shown to select for an increase in cell size (Vasi et al. 1994; Gounand et al. 2016). The observed patterns may also reflect constraints imposed by genome streamlining on the ability of the minimal cell to evolve an adaptive increase in cell size (Mei et al. 2009; DeLong et al. 2010; Chien et al. 2012; Westfall and Levin 2017). With more than 50% of its membrane transport proteins removed, the minimal cell may be unable to sequester the resources needed for constructing and maintaining a larger cell (Chien et al. 2012; Hutchison et al. 2016; Westfall and Levin 2017). However, one might expect that these biosynthetic constraints would be reflected in a reduced rate of adaptation, which was not the case (Fig. 2). In addition, cell size could evolve as a fitness-neutral byproduct of selection on other traits, such as DNA replication rate (Amir 2017). For example, the two strains could have evolved different size trajectories despite similar selection pressures, due to epistatic effects of genome minimization such as those demonstrated using the *ftsZ* E315* mutants (Table S3, Fig. S4). In any case, our findings highlight that cell size, a fundamental feature of biological complexity in multicellular and single-celled organisms alike, evolves in a way that is highly dependent on genomic context.

### Outlook

We documented genes, proteins, and traits that are critical for evolutionary performance in the synthetically constructed *M. mycoides,* a bacterium with the smallest genome of any self-replicating form of life. In its ancestral state, this working approximation of a minimal cell had low fitness, fewer than 500 protein-coding genes, and the highest mutation rate ever recorded in a cellular organism. A single gene disruption could have proven lethal. Instead, the force of natural selection outweighed any deleterious consequences of mutation and drift that could have led populations to extinction. Yet, even in one of the simplest organisms, evolution was still complex. Rapid adaptation involved epistatic interactions and selection on divergent targets, 25% of which encoded for proteins of unknown function. Evolution studies like the ones described here can improve gene characterization and the mapping of regulatory networks, which will inform subsequent rounds of rational engineering and streamlining. In addition, minimal cells can be leveraged to address longstanding problems in biology. Derived from suites of diverse ancestors, they may serve as new models for investigating innovations of early life (Kacar et al. 2017), while providing bottom-up insight into the development of multicellularity (Márquez-Zacarías et al. 2021), the mechanisms of coevolution (Hillesland and Stahl 2010), and the assembly of multi-species consortia and their emergent functions (Goldford et al. 2018).

## METHODS

### Strains and growth conditions

We maintained synthetic *Mycoplasma mycoides* JCVI-syn1.0 and synthetic *M. mycoides* JCVI-syn3B in SP4 medium with KnockOut™ Serum Replacement (Gibco) substituted for fetal bovine serum (Table S4). Cultures were grown in a dark, static growth chamber at 37 °C. The non-minimal JCVI-syn1.0 strain has been described in detail elsewhere (2010). The minimal JVCI-syn3B is identical to the strain synthesized in previous studies (2016) with the following exceptions: JVCI-syn3B possesses a second rRNA operon copy, lacks a gene (MMSYN1_0531) coding for an efflux protein, and has 19 genes that were added back into the minimal genome to render the cell easier to use (Breuer et al. 2019; Pelletier et al. 2021) (Table S5). The strain also contains a landing pad system (*Cre* recombinase and *LoxP*) facilitating genetic manipulation. For competition experiments used to quantify relative fitness, we used a JCVI-syn1.0 strain that expresses mCherry, which allowed us to distinguish it in mixed culture from other strains via flow cytometry (see below).

### Mutation accumulation experiment

#### Overview

Mutation accumulation (MA) experiments are designed to reduce the influence of natural selection through repeated bottlenecks of evolving populations (Lynch et al. 2016). When used with microbial populations, this is typically achieved by transferring single colonies, which have undergone single-cell bottlenecks. Prior to initiating MA experiments, we acclimated JCVI-syn1.0 and JCVI-syn3B to laboratory conditions by maintaining populations in SP4 liquid medium. We took a clone of each acclimated strain to begin the MA experiment. We propagated replicate MA lineages (JCVI-syn1.0: 87 replicates; JCVI-syn3B: 57 replicates) of each strain for 20 to 36 weekly transfers.

#### Number of generations

In order to compare rates of mutation across replicates, we normalized all rates as per-generation values. To calculate the number of generations per transfer in the MA, we grew cells on SP4 agar for one week and diluted a sample of 7^th^ day colonies into 1 mL of phosphate-buffered saline (pH = 7.4). Cells were fixed with 20 μL of 25% glutaraldehyde and stained with 2X SYBR Green, and then counted with a NovoCyte flow cytometer (ACEA Biosciences). We used the dilutions to calculate the number of cells in the original colony, from which we inferred the number of generations (log_2_(*N*), where *N* is the number of cells in the undiluted colony) that must have occurred to reach a colony of that size (Dillon et al. 2017), assuming each colony is formed by a single progenitor cell. Because growth rate and other fitness components can decrease during an MA experiment (Behringer and Hall 2016), we also measured the number of cells per colony during and at the end of the MA, averaging across time points to estimate the total number of generations. We then used the number of generations per transfer to estimate the effective population size (*N*_e_) via the harmonic mean method (Behringer and Hall 2016). Specifically, *N*_e_ was approximated as the harmonic mean of the series (2^0^, 2^1^, 2^2^, …, 2^*f*^), where *f* is equal to the number of generations per transfer inferred from the previous step.

#### Whole-genome sequencing and sequence analysis

We performed DNA extractions from evolved MA lines using a DNeasy UltraClean Microbial Kit (Qiagen) following the manufacturer’s instructions, with the additional step of adding 50 μL of 50 mg/mL lysozyme to improve cell lysis. Genomic DNA was sequenced using Illumina MiSeq sequencing to a depth of ≥35X coverage. Library preparation and DNA sequencing were conducted by the Indiana University Bloomington Center for Genomics and Bioinformatics. Whole-genome sequencing reads were quality controlled using cutadapt (Martin 2011) to trim low-quality base pairs and remove residual adapter sequences. We used breseq with default parameters (Barrick et al. 2014; Deatherage and Barrick 2014) to call mutations using the trimmed reads. We only considered fixed mutations for the MA lines. We checked for mutations that had arisen in experimental ancestor strains prior to evolution. Ancestral mutations were removed from analysis of all evolved lines derived from that strain using gdtools (Barrick et al. 2014; Deatherage and Barrick 2014). We used the sequence data to check for contamination or cross-contamination in evolved lines.

#### Statistical analyses

To compare mutation rate and spectrum between strains, we used two-sample *t*-tests for numerical response variables and two-sample *χ*^2^ tests with continuity correction for comparing proportions. For comparing proportions to theoretical expectations within a strain, we used one-sample *χ*^2^ tests with continuity correction.

### Adaptive evolution

#### Overview

We conducted experiments that allowed bacteria to achieve large population sizes to increase the efficacy of natural selection. This involved serial passaging of cells in liquid cultures with limited bottlenecking at each transfer. For example, in our experiment, the minimum population size was 2-4 × 10^7^ for both JCVI-syn1.0 and JCVI-syn3B. We passaged replicate 3 mL liquid cultures of each strain (*n* = 4 per strain) in 13 mm glass culture tubes *via* 1% v/v serial transfer each day for 300 days in a dark, static incubator held at 37 °C. We calculated the number of generations per day as the log_2_ of the dilution factor, i.e., log_2_(101), the number of binary fissions needed to regenerate the original population size after the 1% v/v transfer (Lenski and Travisano 1994). Based on this, we estimated that the *M*. *mycoides* strains were maintained for 1,997 generations, which based on other experiments, is long enough for the majority of adaptation to occur (Lenski and Travisano 1994; Lang et al. 2011).

#### Measurement of fitness

We measured relative fitness by competing ancestral and evolved strains against a third-party *M. mycoides* JCVI-syn1.0 strain labeled with mCherry (Wiser and Lenski 2015). Each strain was grown in liquid medium to log phase, and then the labeled and unlabeled strains were simultaneously diluted into a mixed culture in fresh medium. The abundances of the labeled and unlabeled strains in the mixed culture were calculated by immediately (*t*_0_) sampling the culture, fixing the cells with 20 μL of cold 25% glutaraldehyde incubated at 4 °C for 20 min, and staining with 2X SYBR Green. After 24 hr of growth (*t_f_*), the mixed culture was sampled again in the same manner. We quantified the abundance of each strain using either our Novocyte flow cytometer (ACEA Biosciences) or an LSR II flow cytometer (BD Biosciences) at Indiana University’s Flow Cytometry Core Facility. The abundance of the strain of interest was calculated from the observed number of SYBR Green-only positive cells while the abundance of the third-party strain was calculated from the number of cells positive for both SYBR Green and mCherry. From these cell counts, we calculated relative fitness as the change in the relative abundance of the strain of interest during the 24 hr period of competitive growth versus the third-party strain. Specifically, the relative fitness versus the mCherry third-party strain *W_C_*

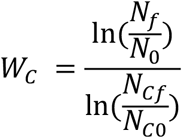

is where *N*_0_ represents the initial abundance of the focal strain, *N_f_* the abundance of the focal strain after 24 hr, and *N*_C*f*_ and *N*_C0_ are final and initial abundances of the mCherry third-party strain, respectively (Wiser and Lenski 2015). We standardized fitness values to be relative to the original *Mm* JCVI-syn1.0 ancestor strain. In other words, we represent the fitness as 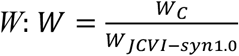, where *W_JCVI–syn1.0_* is the value of *W_C_* for *M. mycoides* JCVI-syn1.0. Axenic control cultures were always run contemporaneously in order account for a proportion of mCherry-positive cells that failed to detectably express mCherry.

#### Whole-genome sequencing and sequence analysis

DNA extraction, sequencing, and bioinformatics followed the same methods as for the MA with a few exceptions. Specifically, each replicate population was sequenced to a depth of ≥ 100X coverage, and polymorphic mutations were included in our analyses. In addition, we calculated *d*_N_/*d*_S_ for evolved lines using the Jukes-Cantor-corrected equation (Yang 2006) as a possible indicator of molecular constraint among the lines. We counted the number of synonymous and nonsynonymous sites using the gdtools module of breseq (Deatherage and Barrick 2014). The observed numbers of synonymous and nonsynonymous substitutions were obtained directly from breseq outputs. Synonymous and nonsynonymous polymorphisms were included in the observed count with probability proportionate to their frequency in mapped reads. We added a pseudocount of 1 synonymous substitution for all calculations (Shpak et al. 2015) because two of the lines had 0 synonymous substitutions.

To identify mutations possibly contributing to adaptation, we looked for genes which had mutations in two or more lines. Mutations in the same gene, arising and increasing in frequency in independent lineages, suggests that mutation’s rise could be driven by positive selection (Johnson et al. 2021). To test this hypothesis, we statistically assessed whether multiply mutated genes had acquired more mutations than would be expected by chance under the assumption that the mutations were neutral (Johnson et al. 2021). To do this, we recorded all of the polymorphic and fixed mutations that were called within genes. Synonymous mutations were excluded. We then simulated placing these mutations at random across all genes. The probability of any given gene receiving any given mutation was proportional to the gene’s length and relativized to the gene’s GC content using the known mutation rates of G:C nucleotides and A:T nucleotides from the mutation accumulation experiment. We repeated this random placement of mutations 100,000 times. In each simulation, we counted the number of mutations received by each gene, with each fixed mutation increasing the count by 1 and each polymorphism counted at a value equal to its frequency. For each multiply-mutated gene from the real adaptation experiment, we measured the proportion of the 100,000 simulations in which the gene received at least as many mutations as were truly observed and called this proportion the *P* value. To correct for the multiple tests (Non-minimal cell: 32; Minimal: 22), we report a Benjamini-Hochberg corrected value *P_adj_* (Benjamini and Hochberg 1995; Benjamini et al. 2009) for which the false discovery rate *α* = 0.05 for the reported significant genes (Table 1).

#### *Generation of* ftsZ *E315* mutant cells*

This process required mutating the bacterial genomes while they were yeast centromeric plasmids (YCPs) followed by genome transplantation of the mutated genomes. The YCPs were mutated using rounds of CRISPR/Cas9 and yeast homologous recombination that is a modification of a method we used previously to mutate *M. mycoides* strains (Kannan et al. 2016).

In the first CRISPR/Cas9 step, the molecule to be mutated was cleaved and the donor DNA comprising sequences from the two flanking genes was recombined with the cut JCVI-syn1.0 or JCVI-syn3B YCP, removing parts of genes of the flanking genes and all of the target gene. The donor DNA had 40 basepair overlaps to both genes flanking the target gene and had a 22 bp *Mycoplasma gallisepticum* 161 CRISPR/Cas9 target sequence with a protospacer adjacent motif (PAM) (5’-GTATAAATACATCCAGGAGtgg-3’) that had no homology elsewhere in JCVI-syn1.0 or JCVI-syn3B. The *M. gallisepticum* sequence put a new PAM in the genome, employed in the second round of CRISPR/Cas9.

The second round of CRISPR/CAS9 cut the JCVI-syn1.0 or JCVI-syn3B YCP at the new *M. gallisepticum* PAM. The cut YCP was then re-circularized using a donor DNA containing the desired point mutation. The mutagenized regions of the YCPs were PCR amplified and the mutation was confirmed by Sanger sequencing. Correctly mutagenized JCVI-syn1.0 or JCVI-syn3B YCPs were then transplanted into *Mycoplasma capricolum* recipient cells as reported previously (Lartigue et al. 2007, 2009; Hutchison et al. 2016). The mutagenized regions of the transplants were PCR amplified and sequenced to confirm the presence of the desired mutations.

#### Microscopy and image analysis

To compare changes in cell size of evolved populations, we centrifuged stationary-phase cultures and resuspended the pellet in 1 mL of phosphate-buffered saline (pH = 7.4). The resuspended cells were fixed by adding 20 μL of cold 25% glutaraldehyde and incubating at 4 °C for 20 min. For microscopic observation, fixed cells were concentrated 50X via centrifugation and resuspension. The centrifugation steps were performed at room temperature for 4 min at 2000 *g*.

For phase contrast microscopy, we pipetted 5 μL of concentrated culture onto an agarose pad (1% agarose in phosphate-buffered saline) and observed cells in the Plan Apo 100X Ph 3 objective lens of a Nikon Eclipse 80i phase contrast microscope. Image data were analyzed using MicrobeJ (Ducret et al. 2016). Cell outlines were drawn using default parameters. Area (number of pixels) within the outline was use as a metric of the area of one face of the spherical cells. Pixel values were converted to μm^2^ using the conversion factor 1 pixel length = 0.0645 μm. Cell diameter was inferred from this area using the assumption of spherical geometry.

Scanning electron microscopy (SEM) was performed at the Indiana University Bloomington Electron Microscopy Center. Fixed cells in PBS were pelleted and resuspended in 100 mM sodium cacodylate buffer (pH = 7.2) with 2 mM calcium chloride and 2% sucrose. We coated 12 mm diameter glass coverslips with 0.1% poly-L lysine for 5 min, after which coverslips were washed with a few drops of double distilled water. Resuspended cells were added to the coverslip surface and allowed to adhere. After 5 min the coverslips were washed twice with 100 mM sodium cacodylate buffer (pH = 7.2) with 2 mM calcium chloride and 2% sucrose. Next, 300 μL of 2% osmium tetroxide in 100 mM sodium cacodylate buffer (pH = 7.2) with 2% sucrose was added to the surface of the coverslips while on ice. After 30 minutes the coverslips were washed with double distilled water. Coverslips were placed in a CPD coverslip holder (Electron Microscopy Sciences, catalog number 70193-01). Samples were dehydrated in a graded ethanol series (30%, 50%, 70%, 90%, 95%) while on ice. At room temperature, coverslips were rinsed three times with 100% ethanol. Each dehydration step lasted 2 min. Critical point drying was done using a Tousimis Samdri 790 critical-point dryer. The dried coverslips were placed on aluminum SEM stubs and sputter coated, using a Safematic CCU-010 with SP-010 Sputter Head, with 45 nm of gold/palladium (80%/20%). We viewed the samples with a FEI Teneo scanning electron microscope at 2.0 kV, 25 pA probe current, and 3.0 mm working distance. The T2 detector was used. We analyzed the SEM image data using ImageJ (Schneider et al. 2012). We used the “straight” and “measure” features combined with image scale metadata to measure the vertical diameters of imaged cells that met the following criteria: cells must be round; cells must not have apparent holes or punctures; cells must be completely within the field of view; cells must have an unambiguous perimeter; there must be no suggestion that a cell is currently or has recently undergone binary fission; cells must ≥ 0.1 μm across. Each image was processed counterclockwise starting from east. The samples were processed in a randomized order.

#### Statistical analyses

We compared relative fitness, *d*_N_/*d*_S_, and cell size between evolved populations using two-sample *t*-tests, and to ancestral values using one-sample *t*-tests. We also used two-sample *t*-tests to compare the relative fitness and cell size of *ftsZ* E315* mutants to the respective ancestral values.

We compared the composition of genes acquiring mutations among the evolved replicate lines by first constructing a gene-by-population matrix. Herein, each row represented an evolved line and each column represented a gene that had acquired at least one mutation among all of the lines. Each cell of the matrix was filled with the sum value of mutations occurring in that gene in that line, where fixed mutations were valued at 1 and polymorphisms were valued equal to the allele frequency. Only essential genes, shared between JCVI-syn1.0 and JCVI-syn3B, were considered. We used a permutational multivariate analysis of variance (PERMANOVA) on the Bray-Curtis distances generated from the gene-by-population matrix to test for the significance of cell type (minimal vs. non-minimal) on the composition of mutations using the adonis function in the R package vegan (Oksanen et al. 2020). For visualization, the Bray-Curtis distances were decomposed into two dimensions using principal coordinate analysis via the cmdscale function.

## ACKNOWLEDGEMENTS

We acknowledge financial support from the US National Science Foundation (DEB-1442246 and 1934554 JTL; MCB-1818344 and 1840320 JIG, LS, and KSW), US Army Research Office Grant (W911NF-14-1-0411 JTL and ML), the National Aeronautics and Space Administration (80NSSC20K0618 JTL), the National Institutes of Health (R35-GM122566-01 and 2017-202 ML), and the Brazilian Agricultural Research Corporation (DMCB). We also acknowledge B. Stein for assistance with scanning electron microscopy (NIH 1S10OD023501-01); J. French, E. Snider, K. McKenzie, B. Lehmkuhl for assistance in the laboratory; and M. Behringer for discussion regarding sequence analyses. Data and code are available at https://github.com/LennonLab/MinimalCell.

