## supplemental for "Evolution of a minimal cell" (from another version of this preprint: version 2, 2021-07-02)

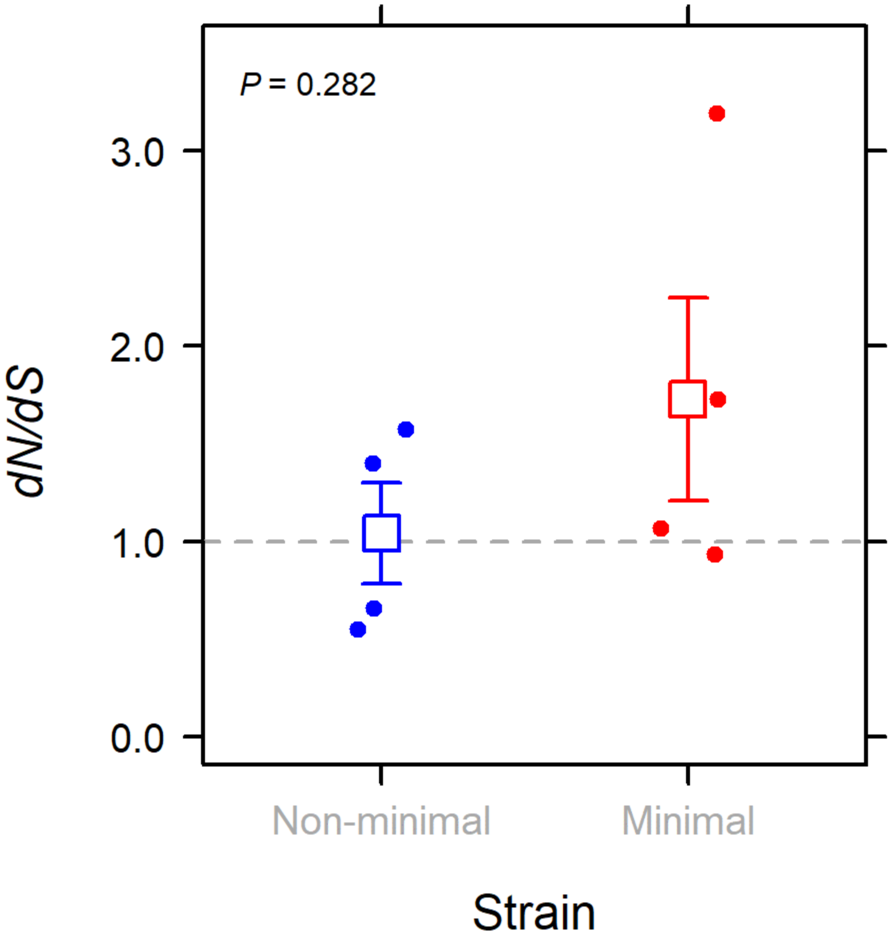


**Figure S1**. The ratio of nonsynonymous to synonymous substitutions in populations of *M. mycoides* after 2000 generations of evolution. The numbers of nonsynonymous and synonymous single nucleotide mutations (SNMs) were counted within each of the evolved replicate populations. The normalized ratio of nonsynonymous to synonymous mutations ($d_N/d_S$) can indicate the relative action of positive or negative selection. Values of $d_N/d_S > 1$ indicate positive selection, while $d_N/d_S < 1$ can indicate the dominance of negative selection and constraint on adaptation. In contrast to expectations, $d_N/d_S$ was not lower for the minimal cell, suggesting that it did not experience more constraint during adaptation. Open boxes represent the mean, error bars represent ± standard error of the mean (SEM), solid symbols represent values for each replicate population. *P*-value corresponds to output from a two-sample *t*-test.

**Table S1.** Mutations observed in *ftsZ* during adaptive evolution across all replicates.

| Strain | Replicate ID | Mutation | Mutation effect | Mutant allele frequency |
|---|---|---|---|---|
| Non-minimal | MM_1 | Q349* (CAA→TAA) | Nonsense | 1 |
| Non-minimal | MM_3 | Q346* (CAA→TAA) | Nonsense | 0.079 |
| Non-minimal | MM_4 | Δ1 bp (597/1158 nt) | Frameshift | 1 |
| Non-minimal | MM_6 | G17C (GGT→TGT) | Missense | 1 |
| Minimal | MM_11 | +CT (1062/1158 nt) | Frameshift | 1 |
| Minimal | MM_11 | V190I (GTT→ATT) | Missense | 1 |
| Minimal | MM_13 | E315* (GAA→TAA) | Nonsense | 1 |

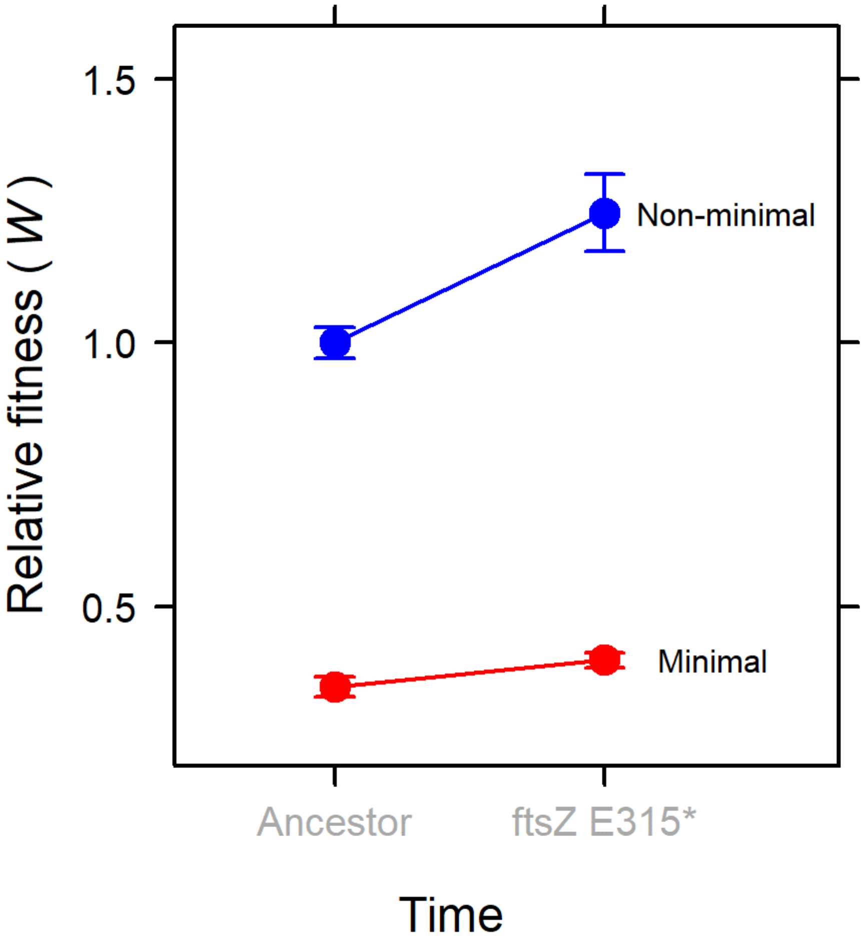


**Figure S2**. Fitness effects of an *ftsZ* mutation on populations of *Mycoplasma mycoides*. We reengineered the nonsense mutation *ftsZ* E315* and quantified its effect on fitness in both the non-minimal and minimal cells. The mutation increased fitness by 25% and 14%, respectively. Data represent means ± SEM.

**Table S2**. Diameter and volume measurements of *M. mycoides* ancestor and evolved cells observed *via* phase contrast microscopy. Values displayed as mean ± SEM.

| | Non-minimal Cell | | Minimal Cell | |
|---|---|---|---|---|
| | Diameter (nm) | Volume ($\mu m^3$) | Diameter (nm) | Volume ($\mu m^3$) |
| Ancestor | 575 ± 28 | 0.123 ± 0.020 | 422 ± 23 | 0.058 ± 0.010 |
| Evolved | 993 ± 53 | 0.577 ± 0.102 | 463 ± 48 | 0.082 ± 0.024 |

**Table S3**. Diameter and volume measurements of *M. mycoides* ancestor, evolved, and *ftsZ* E315* cells observed *via* scanning electron microscopy. Values displayed as mean ± SEM.

| | Non-minimal Cell | | Minimal Cell | |
|---|---|---|---|---|
| | Diameter (nm) | Volume ($\mu m^3$) | Diameter (nm) | Volume ($\mu m^3$) |
| Ancestor | 439 ± 13 | 0.051 ± 0.004 | 305 ± 28 | 0.021 ± 0.002 |
| Evolved | 792 ± 128 | 0.479 ± 0.260 | 375 ± 35 | 0.044 ± 0.012 |
| *ftsZ* E315* | 550 ± 15 | 0.099 ± 0.008 | 246 ± 12 | 0.011 ± 0.001 |

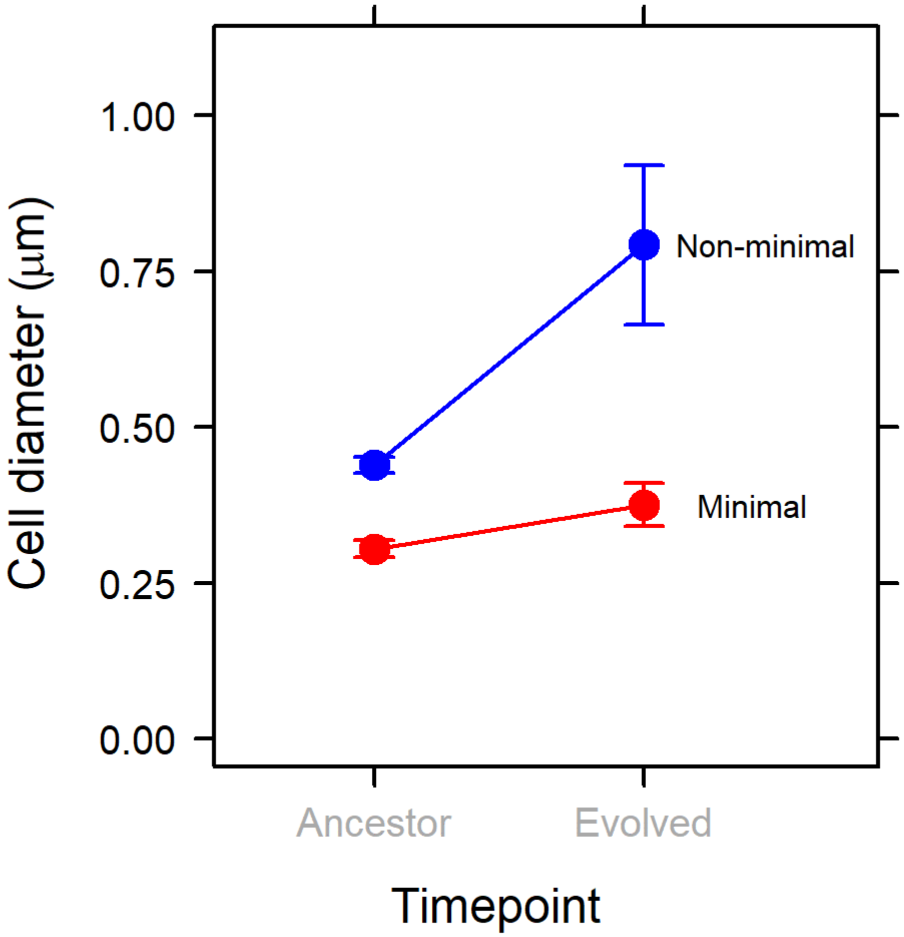


**Figure S3**. Cell size of ancestor and evolved cells from the non-minimal and minimal strains observed using scanning electron microscopy (see Methods). Similar findings are reported in the main text (Fig. 4) using phase microscopy. The evolved phenotypes of the strains diverged: The size of the non-minimal cells increased marginally during evolution ($t_3 = 2.76$, $P = 0.070$), while the size of the minimal cell did not change ($t_3 = 1.01$, $P = 0.459$). Data represent means ± SEM. Error bars for the ancestor represent experimental replicates, while error bars for evolved lines represent evolutionary replicates.

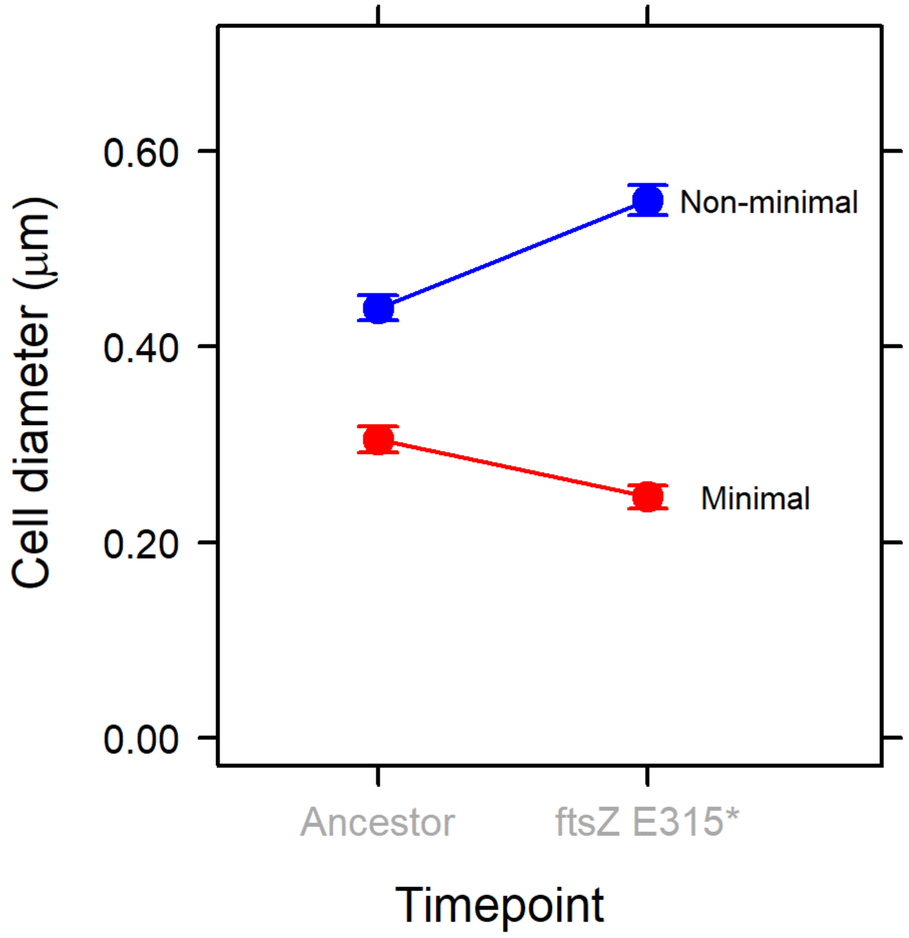


**Figure S4**. Cell size of *ftsZ* E315* mutants compared to the nonmutant for each strain, measured by scanning electron microscopy. The *ftsZ* nonsense mutation caused an increase in wildtype cell size ($P = 8.71 \times 10^{-8}$) but did not increase the minimal cell's size. In fact, the minimal cell *ftsZ* E315* mutant was smaller than the nonmutant ($P = 0.0013$). These observations are in accord with the observation of increased cell size during evolution for the wildtype but not the minimal cell. The data suggest that *ftsZ* mutations could contribute to the mechanism of the divergent cell size trajectory during evolution. Error bars represent ±2 SEM.

**Table S4**. SP4 medium reagents.

| Ingredient | Manufacturer | Quan-tity/L | Unit |
|---|---|---|---|
| Mycoplasma broth base | Hardy Diagnostics | 3.5 | g |
| Bacto tryptone | Fisher BioReagents | 10 | g |
| Bacto peptone | Fisher BioReagents | 5.3 | g |
| Glucose | Sigma-Aldrich | 5 | g |
| Yeastolate | Life Technologies | 2 | g |
| Epure filtered water | | 725 | mL |
| CMRL 1066 (10X) | Corning | 50 | mL |
| 7.5% sodium bicarbonate | Fisher Chemical | 14.6 | mL |
| 200 mM L-glutamine | Fisher Chemical | 5 | mL |
| Yeast extract solution 15% | Gibco | 35 | mL |
| Knockout serum replace-ment | Gibco | 170 | mL |
| Penicillin G | Sigma-Aldrich | 100 | µL |
| Phenol red | Sigma-Aldrich | 0.011105 | g |

**Table S5**. Genes added to *Mycoplasma mycoides* JCVI-syn3B.

| Locus Tag | Annotation |
|---|---|
| MMSYN1_0520 | Conserved hypothetical protein |
| MMSYN1_0521 | Cell division protein sepF |
| MMSYN1_0522 | Cell division protein ftsZ |
| MMSYN1_0527 | Putative metal binding or nucleic acid binding |
| MMSYN1_0538 | Conserved hypothetical protein |
| MMSYN1_0546 | Pseudogene |
| MMSYN1_0548 | tRNA (cytidine(34)-2'-O)-methyltransferase |
| MMSYN1_0549 | Non-canonical purine NTP pyrophosphatase |
| MMSYN1_0592 | Uncharacterized protein |
| MMSYN1_0593 | Uncharacterized protein |
| MMSYN1_0601 | Putative CAAX amino terminal protease, transmembrane |
| MMSYN1_0602 | Pseudogene |
| MMSYN1_0604 | LemA family protein |
| MMSYN1_0605 | Putative phosphate starvation inducible protein (psiF), |
| MMSYN1_0610 | DNA-formamidopyrimidine glycosylase |
| MMSYN1_0622 | Uncharacterized lipoprotein |
| MMSYN1_0623 | Putative adenine glycosylase, methyltransferase, kinase |
| MMSYN1_0930 | 50S ribosomal protein L33 rpmG |
| MMSYN1_0931 | Adenylyl-sulfate kinase met14p |
